## Supplementary information appendix for "Xenosiderophore transporter gene expression and clade-specific filamentation in *Candida auris* killifish infection"

Hugh Gifford^a^, Tina Bedekovic^a^, Nicolas Helmstetter^a^, Jack Gregory^a^, Qinxi Ma^a^, Alexandra C. Brand^a^, Duncan Wilson^a^, Johanna Rhodes^b,c^, Mark Ramsdale^a^, Tetsuhiro Kudoh^d,1^, Rhys A. Farrer*^a,1^

#### Supplementary Results

To examine orthologues across reference genomes for each clade (**Figure S4**), we identified 5,324 orthogroups across clades I-V with outliers *C. haemulonii* and *C. albicans* containing 95.4% of all genes (*n* = 38,345 genes, **Figure S4****A-B**). The use of clade-specific reference genomes identified higher numbers of DEGs (692 *vs* 499 up-regulated and 714 *vs* 523 down-regulated genes) including genes within the accessory genome (55 *vs* 40 up-regulated and 57 *vs* 45 down-regulated, **Figure S4****C**), which we defined as orthogroups of genes not found in every clade. The core genome across all three species and five clades of *C. auris* included 3619 single copy orthologues ($\bar{x}$ = 66.9% *C. auris* genome per clade) and a further 869 single copy orthologues in *C. auris* ($\bar{x}$ = 82.9% *C. auris* genome per clade), of which 133 were unique to *C. auris*. The remaining core genome included 486-585 multiple copy orthologues shared across all species and 110-121 in *C. auris*, of which 17-19 were present in each clade ($\bar{x}$ = 643.2 genes per clade), bringing the total estimated core genome size to 5131.2/5413.2 genes per clade ($\bar{x}$ = 94.8%). A single copy orthologue tree confirmed the basal status of clade V, and synteny plotting was consistent with limited transversions/inversions within *C. auris* outwith clade II (**Figure S4****A**). These findings are consistent with a highly conserved core yeast genome across *Candida* species, especially within the *Metschinikowiaceae* clade, which appears to contain a degree of preserved genome structure.

We observed high correlation between log-fold change values obtained from clade-specific and core (B8441) reference genomes (Pearson’s >= 0.95, *p* <0.001 per comparison). Among DEGs that had no orthologues in the clade I reference genome (**Figure S4****D**), the highest number of clade-unique genes that were differentially expressed was identified in clade IV; other DEGs included genes involved in cell wall, intracellular transport, metabolism, transcription/translation and transmembrane transport (**Table S3**). In terms of siderophore transporters, we additionally identified a clade IV unique gene associated with a siderophore transport GO term that was differentially expressed at both time-points of infection (CJJ09_005327). Overall, the *in vivo* expression profile across five *C. auris* clades indicated a potential role for the expanded gene family of siderophore transporters during *C. auris* in-host survival and pathogenicity, which was up-regulated compared to growth in nutrient-rich laboratory media.

Comparing transcript levels across *MTL* genes in all five clades, we observed poorer detection of *MTL****α*** transcripts when aligned to the clade I reference genome, as expected (**Figure S6****A**). Direct transcript-level testing, with correction for multiple testing to minimise false positives, indicated that only up-regulation of *PIKA* and *PGA30* (48 HPI) and down-regulation of *THI4* (24 HPI) and *MDR1* (at both time-points) were significantly different across the set of 10 DEGs shared across all comparisons (**Figure S6****B**).

We used the clade I reference genome to annotate functional domains and discover enriched gene families, revealing enrichment of up-regulated hypha-regulated cell wall GPI-anchored proteins in more virulent clades (**Figure 6**), driven by three pairs of orthologues of *IFF4*, *HYR3* and *RBR3* (**Table S3**). Four secretory lipase *LIP1* paralogues drove an enrichment of the related PFAM domain for clade I *vs* III at 24 HPI. When comparing clade I and IV infection to clade II and 48 HPI, we also observed up-regulated nutrient transporters including a 7 transmembrane domain putative ferric reductase/iron importer, *CFL4*, as well as ferric reductase, *FRE3*, and siderophore transporter orthologues *SIT1_1948*, *SIT1_2241* and *SIT1_4097*. Cell wall components, non-mating *MTL* locus genes such as *PIKA*, and genes involved in iron and siderophore transport are therefore potential modulators of strain-specific differences in virulence.

### Supplementary Methods

**Ethical statement:** All animal experiments were performed under Project Licence PP4402521 in compliance with University of Exeter Animal Welfare Ethical Review Board regulations and MRC Centre for Medical Mycology Project Review Board, and are reported in line with “Animal Research: Reporting *in vivo* Experiments” (ARRIVE 2.0) Guidelines^1^.

***Aphanius dispar* care, monitoring, husbandry and procedures:** Wild-type adult AK were kept at the Aquatic Resources Facility, Exeter, in two 28 °C incubation tanks, with 12-hour day and night light cycles and artificial enrichment foliage. Embryos were collected using three plastic netted collection chambers per tank with transfer of all enrichment into tanks, and placed overnight from 16:00 to 09:15 in dark conditions. Chambers were then drained, cleaned and strained with 35 parts per thousand artificial seawater (ASW) to isolate all embryos. Subsequent ASW was sterile suction filtered *via* 0.2 μm pore Nalgene Rapid-Flow (Thermo Scientific, UK). Embryos were inspected for viability in terms of an intact blastocyst without discolouration by stereo dissection microscope (Optech, UK) and incubated at 30 °C (Memmert, Germany) for 72 h with day and night cycles with daily ASW changes and removal of dead, unhealthy or underdeveloped embryos, with a maximum of 30 embryos per standard size 100 mm x 15 mm petri dish (Greiner Bio-One, Austria). Excluded embryos were killed or disposed of either by submission in Virkon solution (Sigma-Aldrich, UK) or freezing at -80 °C.

**Experimental procedures:** Injection needles were prepared using borosilicate glass capillaries (Harvard apparatus, USA) and a heated pulling system (heater level >62, PC-10, Narishige, UK). We used tweezers to snap the tip to an estimated 1 mm graticule-measured aperture of 10-20 μm, and sterilised needles before use with 253.7 nm UV light over 15 min (Environmental Validation Solutions, UK). Injection bays were made with microwaved 1% agarose in 1 g (Sigma, USA) per 100 mL ASW for up to 24 embryos in 100 mm petri dishes using custom-made molds in an inverted 60 mm petri dish. Embryos were randomly chosen from each dish and the sequence of injection for each biological repeat was randomly allocated by computer. For microinjection, *C. auris* yeasts were counted by haemocytometer and added to 25 μL sterile filtered phenol red solution 0.5% (Sigma, UK) to a concentration of 0.5x10^8^ per mL in 100 μL. Injection needles were loaded with 5 μL injection solution using 20 μL Microloader EpT.I.P.S. (Eppendorf, UK) immediately before injection of up to 30 embryos before re-loading. Injections were performed with 400-500 hPa injection pressure and 7-40 hPa compensation pressure over 100-200 ms with the FemtoJet 4i (Eppendorf UK) using a micromanipulator (World Precision Instruments, UK) and a Microtec HM-3 stereo dissection microscope (Optech, UK). In order to deliver a dose of approx. 500 yeasts in 10 nL solution, an injection bolus was administered to an estimated diameter of 0.2 – 0.3 mm (10-15% of total embryo diameter) *via* a single chorion puncture. Any embryos that failed to meet these criteria for injection were excluded. Control *in vitro* inoculations were microinjected as above into 1 mL YPD in 1.5 mL Eppendorf tubes before incubation without shaking at 37 °C for 24 h contemporaneously with embryos as below.

**Study design and sample sizes:** We used an unblinded prospective virulence study comparing six groups of AK embryo yolk-sac microinjection, including five different clades of *C. auris*, and one control sham injection as the negative control. We used >= 23 embryos per condition based on a power calculation of $\sigma$ = 0.25, significance = 0.05, power = 0.9, and $\delta$ effect size of 25% mortality (two-sided t-test). After excluding embryos that were unhealthy or did not complete successful microinjection, We included *n* = 10-13 embryos per group per experiment (total *n* = 34-37 embryos per group). Each embryo was considered to be an experimental unit (*n* = 70-72 embryos per experiment, total *n* = 213 embryos). For estimation of *C. auris* in-host growth by homogenisation and plating of colony forming units, three embryos per condition per time-point across two (0 and 24 HPI) or three (48 HPI) experiments were used (total *n* = 126). For time lapse microscopy demonstration of *A. dispar* yolk-sac collapse and death, representative image sets were chosen from a pilot experiment using a reduced dosage (1/10) of *C. auris* (*n* = 4). For histological demonstration of *C. auris* morphology, one embryo per condition at 24 HPI and 48 HPI each were used in one experiment, and seven embryos per condition plus three embryos without injection in a second experiment (*n* = 57). RNA was extracted from three embryos per condition per time-point (0, 24 and 48 HPI, *n* = 54) plus three embryos per time-point without any injection (*n* = 9, total *n* = 63) as a control for the impact of microinjection.

***C. auris* strains and growth:** We selected five *C. auris* strains to represent each major clade (**?@tbl-Ch3strains**). Culture stocks were kept in -80 °C in Yeast extract Peptone Dextrose (YPD) broth (Sigma-Aldrich, UK) containing 25% glycerol (ThermoFisher, UK) and plated onto YPD-agar (Sigma-Aldrich) for 48 h before innoculation and overnight growth in 10 mL YPD broth at 30 °C in a shaking incubator at 200 RPM (Multitron, Infors HT, UK). Five mL from each overnight culture was then centrifuged at 5000 g for 2 min before triple washing in 1 mL autoclaved deionised water (DIW, Milli-Q, Merck, USA) to prevent aggregation^2^, then re-suspended again in 200 μL DIW. Decontamination of surfaces was conducted with both mopping and cleaning wipes in 70% Ethanol (Sigma-Aldrich) or 10% Chemgene HLD_4_L (Medimark Scientific, UK) as per manufacturer’s instructions. For growth curves, *C. auris* strains were grown overnight in YPD in 4 replicates. Overnight cultures were inoculated into YPD to a final OD_600_ of 0.02 in a 96-well plate. OD_600_ was read over 25 hourly cycles at 37 °C with 300 s shaking at 2.5 mm amplitude on an Infinite 200 PRO plate reader (Tecan, Switzerland).

**Outcome measurement:** Individual embryos were kept in 1 mL sterile filtered ASW per embryo in a 48 well flat bottom plate (Corning, USA) sealed with Parafilm “M” (Bemis Company, USA) at 37 °C with passive humidification using a 500 mL open glass beaker of water. Embryos were monitored for presence of heart rate at 24-hourly intervals up to 7 d by stereo dissection microscope. Initial time of injection was subtracted from each time-point to minimise bias between groups. Kaplan-Meier survival curves were plotted with Log-Rank testing and Benjamini-Hochberg (BH) correction^3^ using survival v.3.5.8 and survminer v.0.4.9 packages in R v.4.4.0. Colony forming units (CFUs) were calculated by homogenisation of embryos at 0, 24, and 48 HPI by autoclaved plastic micropestle in a 1.5 mL conical tube (Eppendorf, UK) in 100 μL 10,000 U/mL penicillin/streptomycin (P/S) solution (Gibco, UK). Dilutions were made with P/S solution and DIW in 1/10-10,000 and counts were adjusted according to dilution after plating onto YPD agar plates and read after 72 h incubation at 30 °C. Statistical testing between CFU numbers was performed with Wilcoxon testing and BH correction.

**Imaging:** For direct microscopy, AK were kept in 300 μL ASW containing 0.04% Tricaine anaesthesia (Sigma-Aldrich, UK) in an optical adhesive and parafilm-sealed 96-well plate. These were then monitored for heartbeat by 5 bright field 2X or 4X images at pre-focused z slices taken every 2 h by Acquifer microscopy (Bruker, Germany) over 72 h. All images were processed in ImageJ 1.53a (NIH, USA). Microscopy of YPD-agar grown colonies was performed without staining in DIW.

**Histology:** Sectioning and staining was performed by Microtechnical Services, Exeter, UK. Embryos were fixed in 1 mL 10% neutral-buffered formalin (Sigma-Aldrich, UK) for >24 h, followed by immersion in 4% phenol 10 mL glycerine BP made up to 100 mL with distilled water^4^, sectioned at 4 μm without orientation, and stained with Harris Haematoxylin & 1% Alcoholic Eosin on Leica Autostainer. H&E stained histological sections were imaged on an Olympus IX83 microscope coupled with an Olympus DP23 colour camera using an Olympus UPlanXApo x40/0.95 objective controlled by cellSens software v.3.2. Brightfield microscopy was performed with an EVOS M5000 microscope (ThermoFisher, UK).

**RNA extraction and sequencing:** Total RNA was extracted using the Monarch® Total RNA Miniprep Kit (NEB, USA). Embryos were flash frozen in liquid nitrogen, re-suspended in DNA/RNA Protection Reagent and disrupted with an autoclaved plastic micropestle. Bead beating was performed in a FastPrep 24 homogeniser (MP Biomedicals) with 0.5 mm zirconia/silica beads for 8 cycles of 35 s at 6 m/s plus 45 s on ice. Lysate was recovered and incubated for 5 min at 55 °C with proteinase K before addition of lysis buffer and RNA purification following manufacturer’s instructions. Total RNA samples were quantified using the Qubit 4.0 Fluorometer (Invitrogen, USA) and integrity was checked with the Tapestation 4200 (Agilent, USA). Library preparation and sequencing were performed by the Exeter Sequencing Service. Samples were normalised and prepared using the NEBNext Ultra II Directional RNA Library Prep - Poly(A) mRNA Magnetic Isolation Module following the standard protocol. Libraries were sequenced on a Novaseq 6000 sequencer with 150bp paired-end reads. Reads were trimmed with fastp^5^ v.0.23.1 to remove reads <75 bases and trim based from the 3’ end with q-score <22. Quality control was performed with FastQC (https://github.com/s-andrews/FastQC) v.0.11.9 and MultiQC^6^ v.1.6.

**Differential gene expression:** We aligned sequences ($\bar{x}$ = 52.9 m reads per replicate, range 41.1-112.2 m) using the Trinity pipeline^7^ v.2.15.1 to the *C. auris* clade I B8441 reference genome (GCA_002759435.2)^8^ downloaded from GenBank (mean aligned reads in infected embryos 3.83 m, 0.81-13.1%). We further aligned clade-specific reference genomes for each strain (**?@tbl-Ch3strains**), including clade II (GCA_003013715.2)^9^, III (GCA_002775015.1)^10^, IV (GCA_008275145.1)^10^ and V (GCA_016809505.1)^11–13^. We aligned conditions containing *A. dispar* to the *de novo* GJEY01 adult gill transcriptome assembly^14^ downloaded from GenBank, for which we detected expression over 27,481 potential genes. Alignment was performed with Bowtie2^15^ v.2.5.1, transcript estimation with RNA Seq by Expectation Maximisation (RSEM)^16^ v.1.3.3, differential expression with EdgeR^17^ v.3.40.0, Samtools^18^ v.1.17 and limma^19^ v.3.54.0 using a false discovery rate (FDR) p-value cut-off of 0.001 and a log fold change cut-off of +/-2.0. Principal components were calculated and plotted using the prcomp function in Rv4.4.0 and ggfortify v.0.4.17 R package for fragments per kilobase per million, and correlations plots from counts per million as per the default output of the Trinity pipeline. Log fold change was plotted with ComplexHeatmap^20^. In creating volcano plots, we did not display a hypothetical host protein without annotations (HP_DN122662) with a log-fold change of -11.11 and a false discovery rate of 30.57 when comparing sham injection *vs* no injection. Additionally, we did not display four *C. auris* transcripts with infinite calculated FDR comparing in-host to *in vitro* DEG at 24 HPI, including *TNA1_4893* and *NGT1_2864* (up-regulated in-host) and *IFF4_4451* and *ALS4_4112* (down-regulated *in vitro*).

**Annotation and enrichment:** Gene ontology terms^21,22^ were assigned using DIAMOND^23^ v.2.1.8 *via* the DIAMOND2GO pipeline^24^ for each of the *C. auris* reference genomes. KEGG terms^25^ were assigned using the BlastKOALA online platform using the eukaryote database^26^. Protein FAMily (PFAM) domains^27^ were identified using the PFAM-A.hmm database (https://ftp.ebi.ac.uk/pub/databases/Pfam/current_release/Pfam-A.full.gz) and HMMER and HMM Scan^28^ with a cut-off of 1e^-5^. Further domain-specific identification was performed with SignalP 5.0^29^, DeepTMHMM 1.0^30^ and NetGPI 1.1^31^. Enrichment analysis for DEG subsets was calculated using the Fisher’s Exact test, adjusted using the Benjamini-Hochberg method per set of up-regulated or down-regulated features in each comparison as a conservative estimate of significant gene set enrichment, with a false discovery rate cutoff of *p* = 0.001. Annotation tables were checked with additional information from available literature, supplemented by details and web-scraped orthologue names from *Candida* Genome Database^32^ and Blastp searches^33^. Annotations for the GJEY01 genome were also performed using Diamond2GO, BlastKoala and HMMScan as above and checked against *A. dispar* annotations and gene names obtained directly from the GJEY01 transcriptome makers^14^. For plotting, lists of GO terms were identified (and for *A. dispar* enrichment, reduced by removal of redundant terms) by REVIGO^34^.

**qRT-PCR:** Quantitative PCRs were performed on three replicates of cDNA reverse transcribed from RNA. Equal quantities of RNA from each sample were reverse transcribed using MMLV reverse transcriptase and oligo dT primers (Promega, UK). qPCRs were carried out using PowerTrack SYBR Master mix (Applied Biosystems, UK) with cDNA as template on a QuantStudio™ 7 Pro qPCR platform (Applied Biosystems, UK). Fold change in gene expression level was calculated using the 2−∆∆Ct method with normalisation to the housekeeping genes *B2MG* (beta-2-microglobulin-like, DN135862|c3_g2_i2) in *A. dispar* and *ACT1* (B9J08_000486) in *C. auris*. Statistics was performed using GraphPad Prism v.10.2.1.

**Orthologue assignment:** To identify accessory genes across *C. auris* clades, we used the Synima pipeline^35^ with Orthofinder v2.5.5^36^ with additional reference genomes for *C. albicans* (SC5314_GCA_000182965.3) and *C. haemulonii* (B11899_ASM1933202v1). We also incorporated a very recent version of the *C. auris* B8441 assembly (v3) to arrange contigs and illustrate re-arrangements, with broad syntenic concordance; it contains 5 more genes than v2, for which 27 (0.5%) and 16 (0.3%) genes were unique to each genome, respectively. Multiple alignment of single copy orthologues as defined by Orthofinder was performed with Muscle v5.1^37^ and a phylogeny was made with FastTree v2.1.11^38^ using default settings.

**RNA-seq meta-analysis:** NCBI SRA was searched for “*Candida auris*” on 16th April 2024 and downloaded as metadata *via* the “Send to: Run Selector” function. Each project accession number for runs containing RNA was searched against available published literature to identify publications, noting subject of study, clades/strains, growth media, comparative conditions, temperature and time scales used. Calculation of FPKM was performed and analysed as above using the set of single copy orthologues to demonstrate variance *via* PCA with ggfortify and prcomp packages in R and to calculate an unweighted co-expression Pearson’s correlation (cut-off of 0.85) for *SIT1* genes with and without the additional data from the RNA-Seq meta-analysis.

***SIT1* analysis:** The protein transcript for *C. albicans* *SIT1* was Blastp searched against the draft pangenome database as above. We visualised the structure using Alphafold v3^39^ and ChimeraX v1.8^40^. Clade I-V *C. auris* and *C. haemulonii* *SIT1* genes were aligned using Muscle as above before using RAXML-NG v1.2.2 with 1000 bootstraps and the LG4X model^41^, which was visualised using ggtree v3.12.0. For panfungal *SIT1* analysis, we used the fungidb portal for Blastp using the *C. albicans* *SIT1* against all available fungal data (with no hits within oomyceta), including 274 reference genomes representing 197 species and a cut-off of 1e^-20^. Annotations for each protein transcript were performed as above to obtain GO terms, KEGG pathways, PFAM domains, GPI-anchors, SignalP domains, and transmembrane domains, while a phylogeny for *SIT1* similar sequences across all fungal references was constructed as above following multiple alignment with Clustal Omega v1.2.4^42^.

**Data availability:** Raw reads have been deposited *via* NCBI/GEO *via* accession [GSE277854](https://www.ncbi.nlm.nih.gov/geo/query/acc.cgi?acc=GSE277854). All datasets and code required to make figures are available *via* Github (<https://github.com/hughgifford/Arabian_Killifish_C_auris_2024>).

**Supplementary Figures**

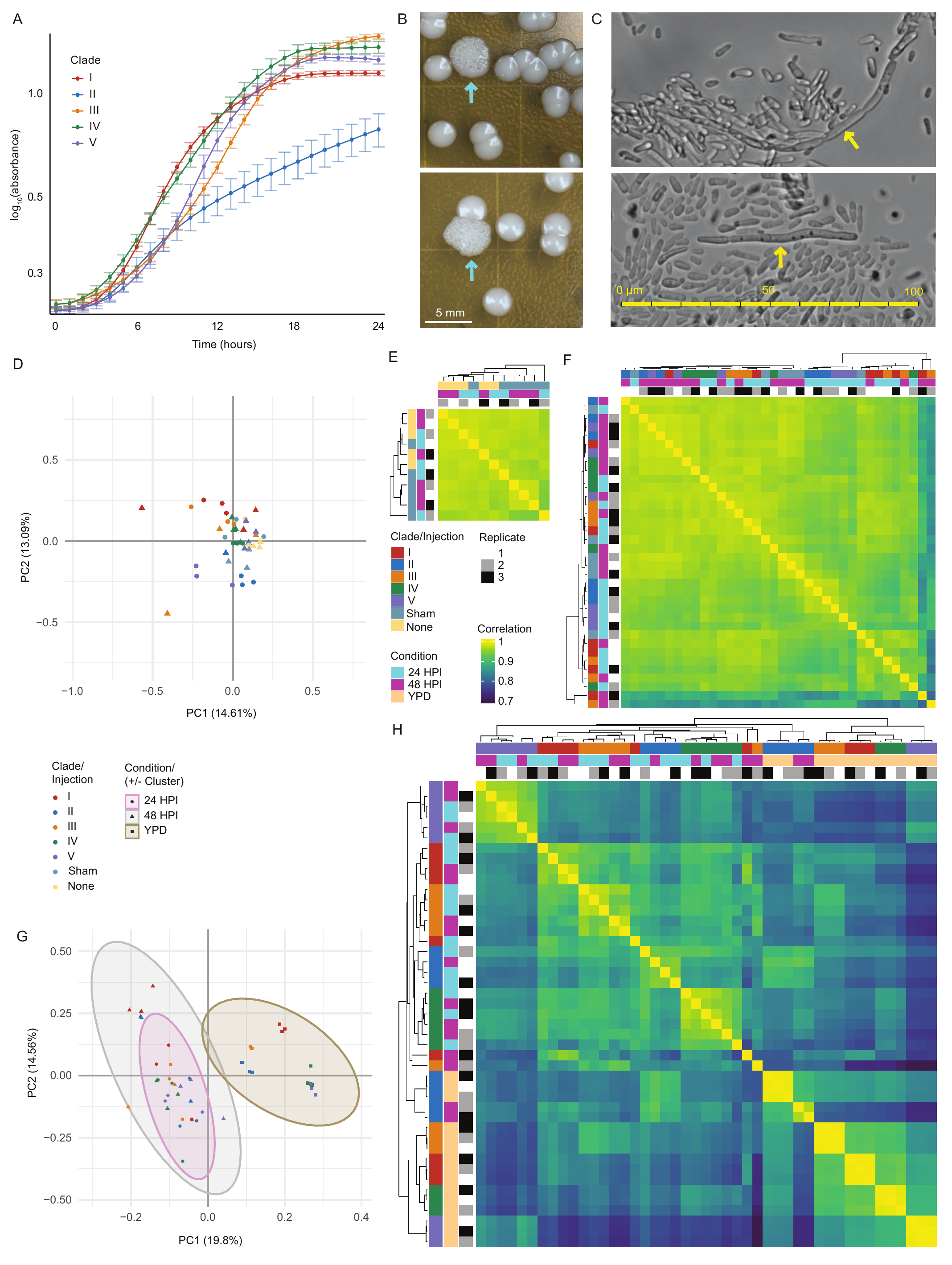

Figure S1: Profiles of *C. auris* growth, morphotype, morphology, and gene expression. **(A)** Growth curves for five representative strains of *C. auris* over 24 h *in vitro*. **(B)** Rough colonies demonstrated on YPD-agar after recovery from AK infection in clade V only (blue arrows). **(C)** Filamentous forms (yellow arrows) isolated from rough colonies and imaged on light microscopy in distilled de-ionised water. **(D)** Principle components analysis (PCA) of fragments per kilobase per million (FPKM) gene expression for AK host transcription. **(E)** Correlation matrix for individual replicates’ log_2_-transformed transcript counts per million for AK responses to sham injection and no injection **(F)** Correlation matrix as above for AK responses to *C. auris* infection and sham injection. **(G)** PCA of FPKM gene expression for *C. auris* pathogen transcription. **(H)** Correlation matrix as above for *C. auris* gene expression *in vivo* and *in vivo* across five clades.

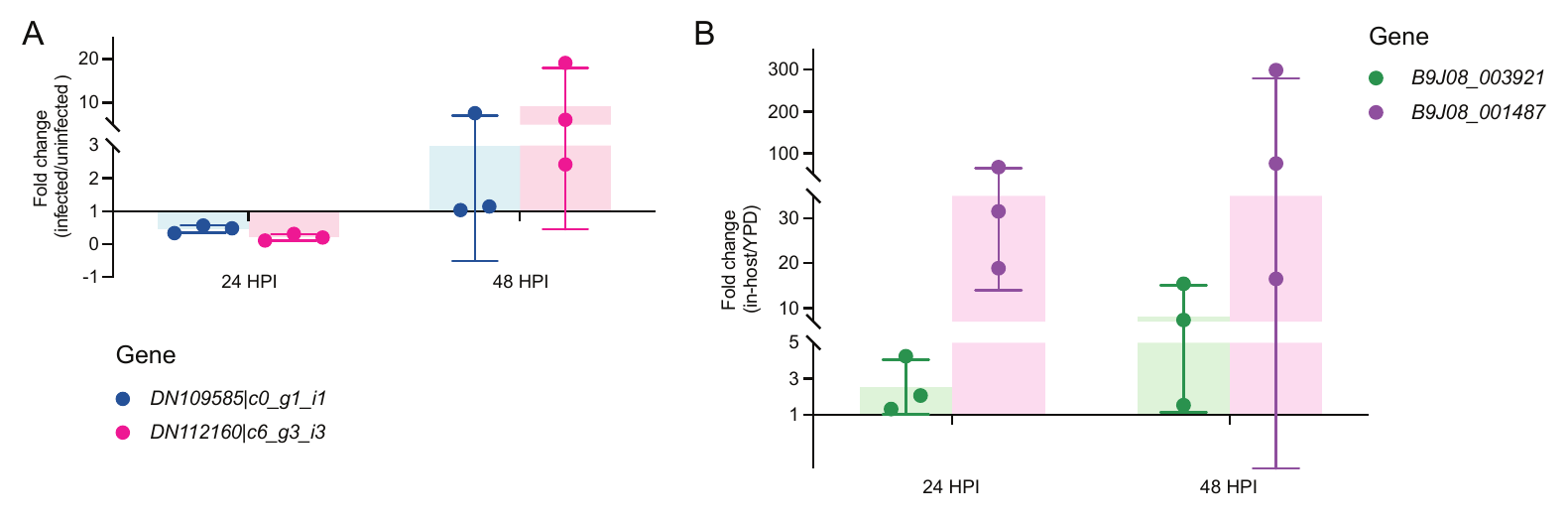

Figure S2: Comparing fold-change by qPCR for RNA recovered from clade IV infected embryos to confirm host and pathogen findings. **(A)** Fold change for two *HMOX* genes, DN109585|c0_g1_i1 and DN112160|c6_g3_i3. Bars represent mean of three replicates; error bars represent standard deviation. **(B)** Fold change for two *XTC* genes. Bars represent mean of three replicates; error bars represent standard deviation.

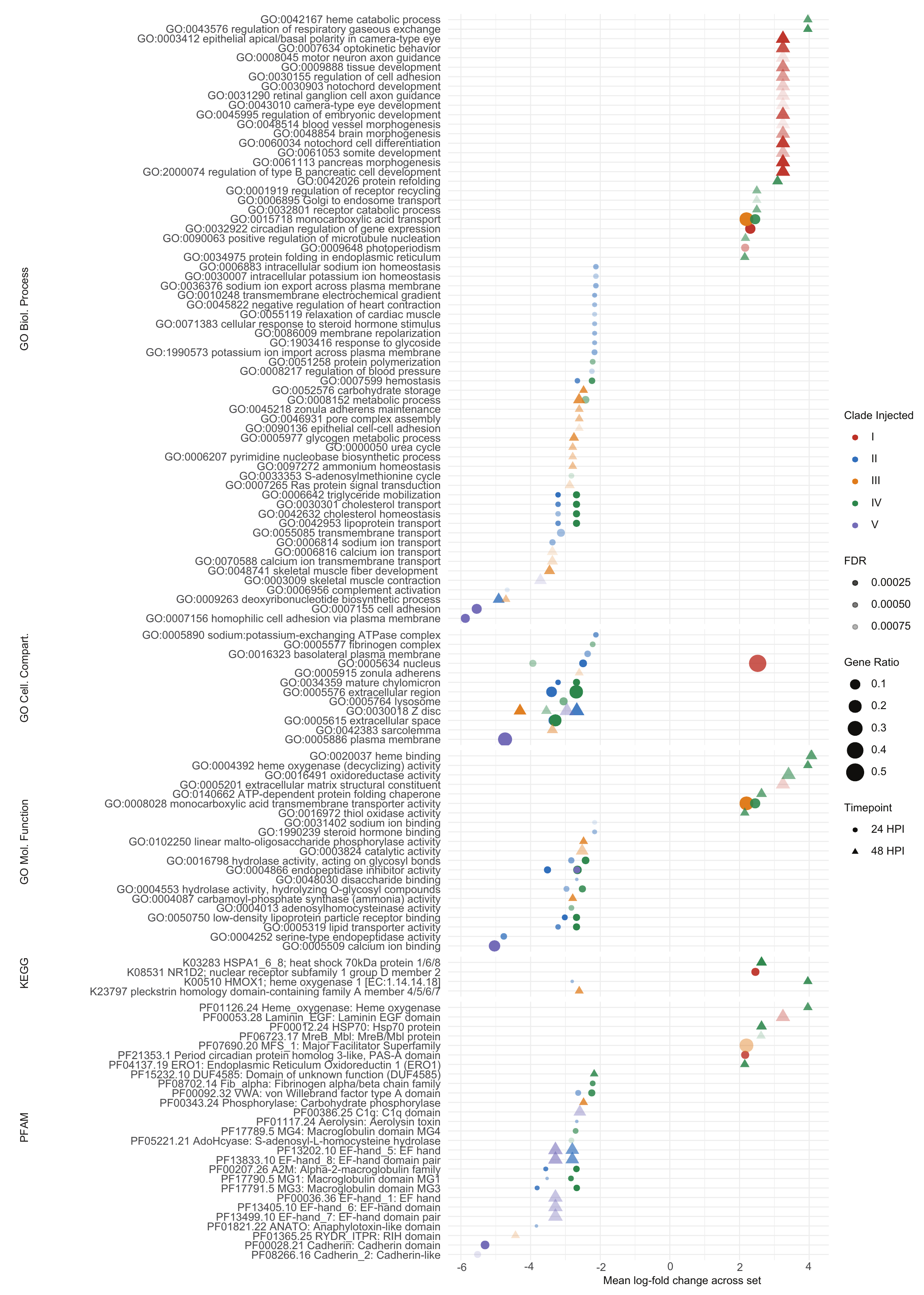

Figure S3: *A. dispar* gene ontology, pathway and domain enrichment during infection *vs* sham control: The GJEY01 *A. dispar* transcripts were annotated for Gene Ontology (GO) terms, Kyoto Encyclopaedia of Genes and Genome (KEGG) pathways and PFAM domains. Enrichment testing was performed with Fisher’s exact test with Benjamini-Hochberg (BH) correction for multiple testing with a false discovery rate (FDR) cut-off of 0.001. Mean log-fold change and ratio of genes in that set was calculated for genes in each set possessing each feature.

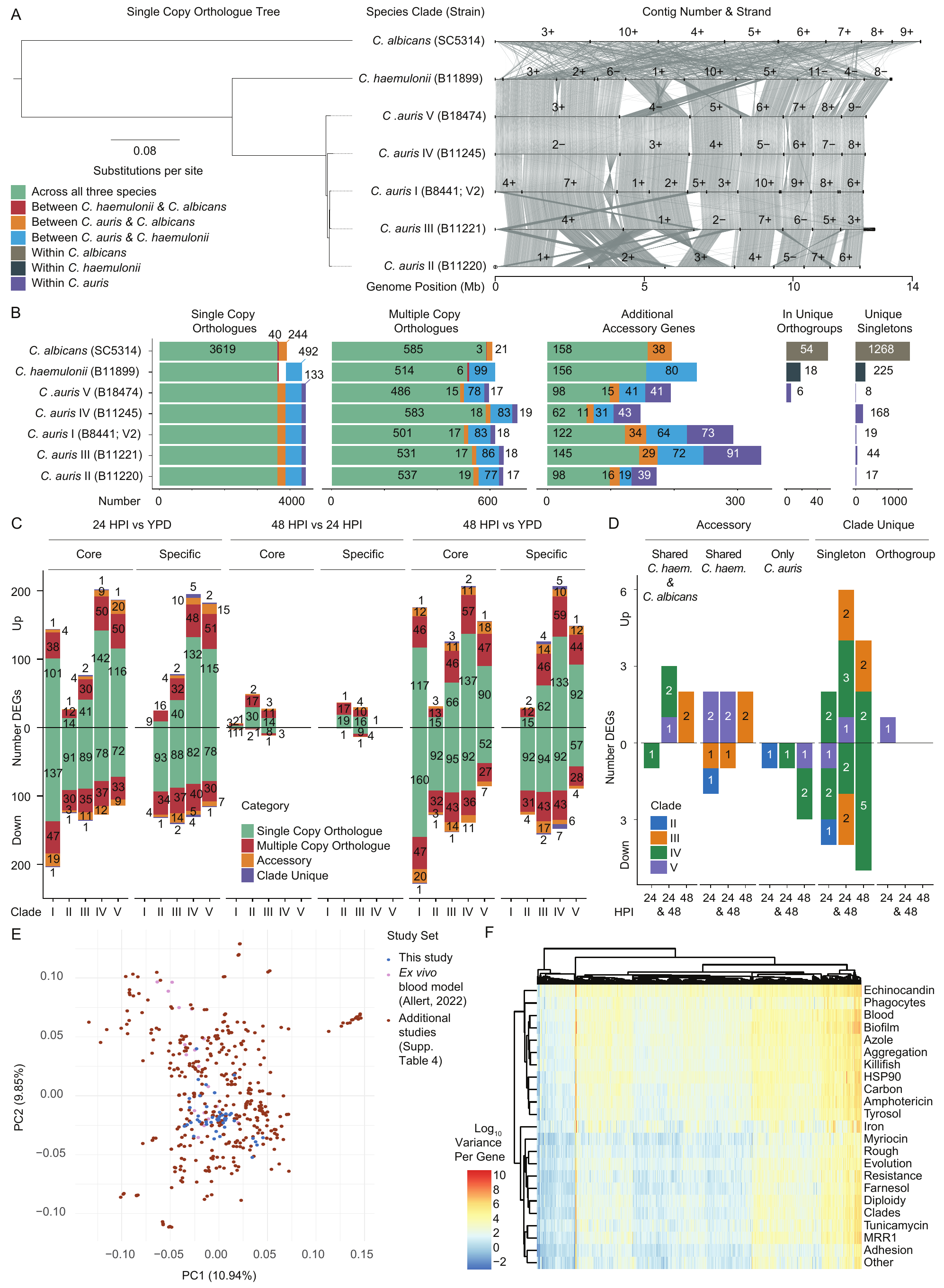

Figure S4: Contextualising differential gene expression across orthogroups and broader RNA-seq datasets. **(A)** Single copy orthologue phylogeny synteny. **(B)** Shared/unique orthologue counts. **(C)** Differential gene expression for *in vivo vs in vitro* comparisons, comparing numbers of DEGs identified when using a single core reference genome (B8441, clade I) or a clade-specific reference genome for each individual clade. **(D)** Differential expression of specific genes not present in core reference genome. **(E)** Per-locus variance of single copy orthologues from RNA-seq meta-analysis of 36 public datasets for *C. auris* transcriptome expression. **(F)** Hierarchical clustering of per-gene variance, calculated for all B8441 core genes for each study.

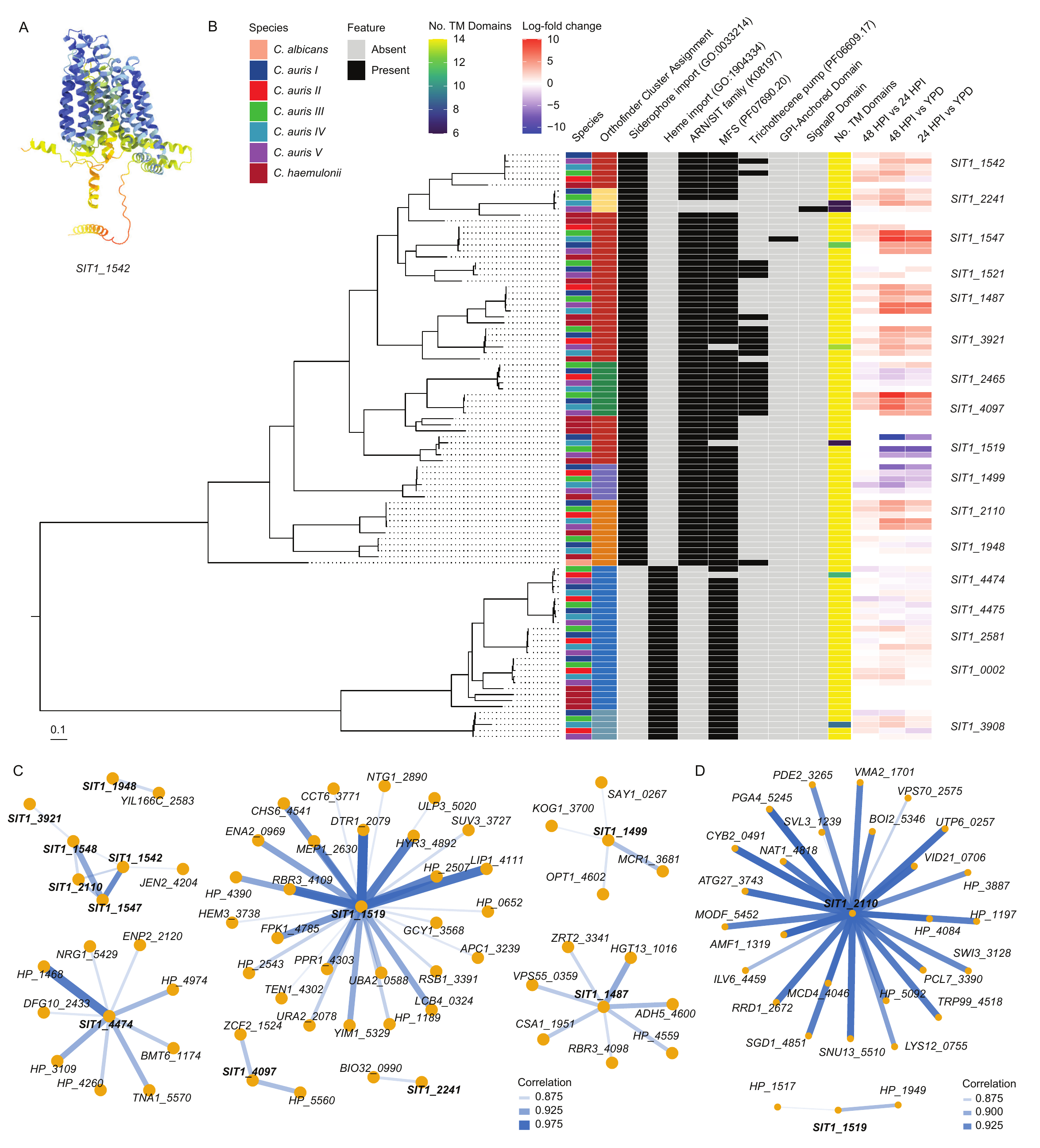

Figure S5: Xenosiderophore Transporter Analysis: **(A)** AlphaFold 3 predicted protein folding for peptide sequences for *SIT1* 1542, illustrating the conserved 14-transmembrane domain similarity across the families. **(B)** RAXML phylogeny using 1000 bootstraps and LG4X model of Muscle-aligned protein sequences for orthologues of *C. albicans* Siderophore Transport 1 *SIT1* gene. Scale bar indicates number of substitutions per site. Orthofinder cluster assignment notes which grouping each gene was assigned to by our Synima/Orthofinder pipeline. **(C)** Pearson’s correlation co-expression of associated gene expression involving siderophore transport candidate genes. **(D)** Pearson’s correlation co-expression across available RNA-seq data from meta-analysis of 35 studies (**Table S4**), demonstrating a different set of transcriptional co-expression indicated by external experimental data.

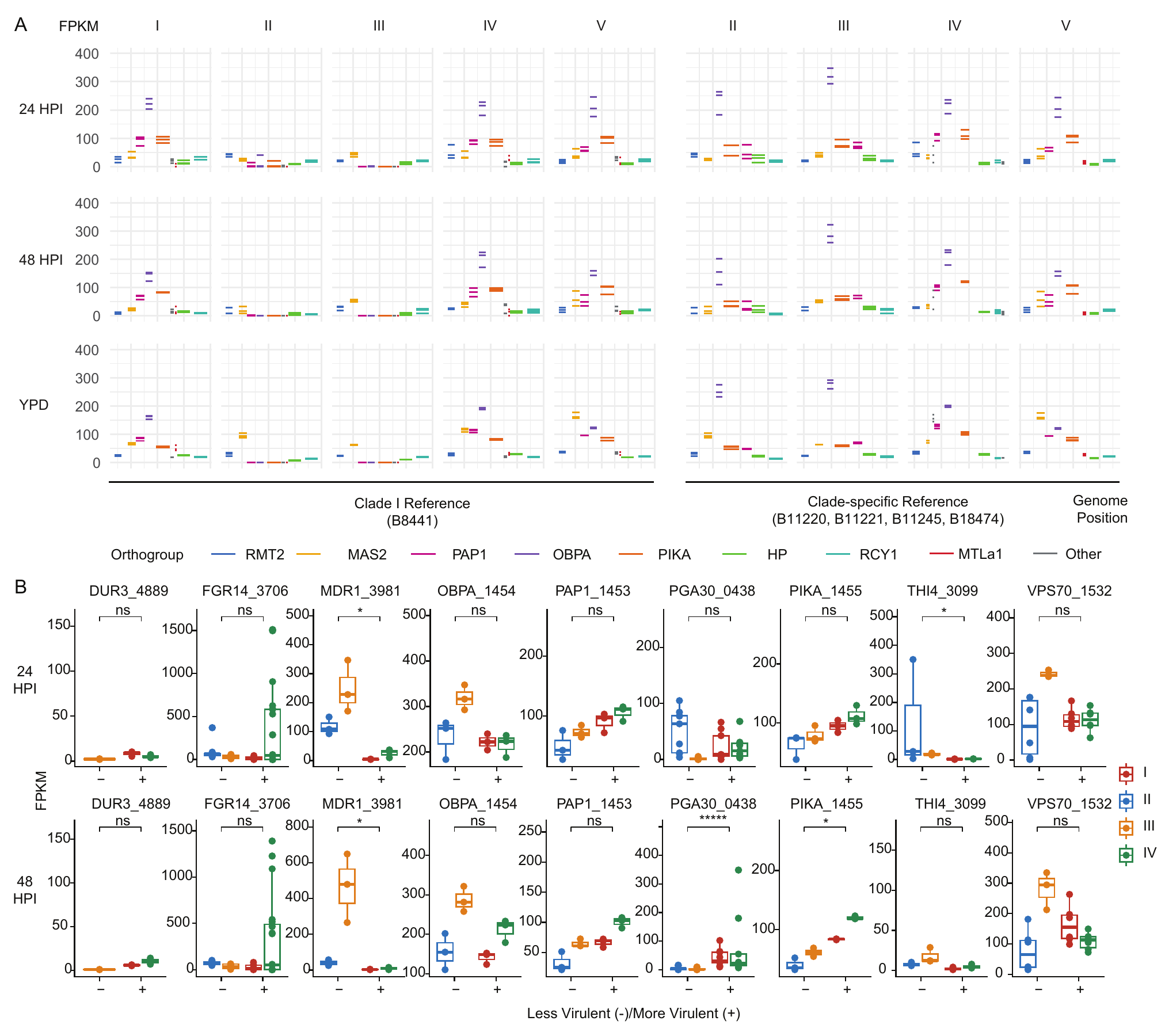

Figure S6: *C. auris* Mating-Type Locus Gene Expression: **(A)** Transcripts (fragments per kilobase per million, FPKM) for orthogroups aligned to the same reference genome (left-hand panels) and clade-specific reference genomes (right-hand panels), indicating similar transcript levels between clades for e.g. *PIKA*, *OBPA* and *PAP1* non-mating genes when aligned to clade-specific reference genomes. **(B)** Comparing individual transcript levels for each orthogroup in more *vs* less virulent clades *via* Wilcoxon signed-rank test with Bonferroni correction to minimise false positives, demonstrating that only down-regulation of *THI4* (at 24 HPI), *MRD1* (at both time-points), and up-regulation of *PGA30* and *PIKA* were significantly different. Significance levels are as follows: * *p* ≤0.05, ** ≤0.01, *** ≤0.001, **** ≤0.0001, ***** ≤0.00001.

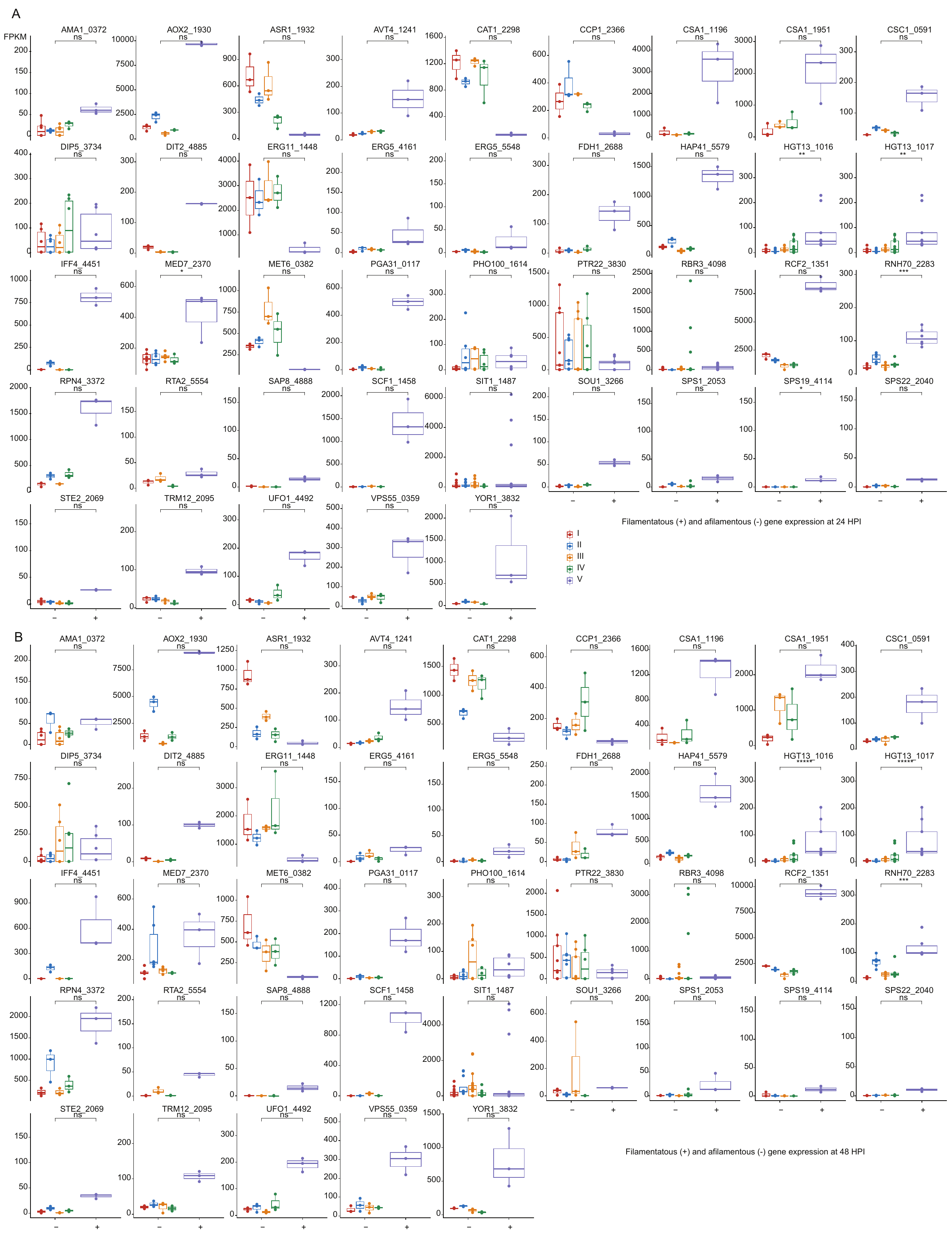

Figure S7: Comparing individual transcript levels for each orthogroup in filamentous clade V *vs* all other clades *via* Wilcoxon signed-rank test with Bonferroni correction to minimise false positives, demonstrating at 24 HPI **(A)** and 48 HPI **(B)**, excluding any genes that were assigned to be ‘hypothetical proteins’ only, up-regulation of two *HGT13* orthologues and *RNH70* were significantly different at both time-points, with up-regulation of *MED7* and *SPS19* at 24 HPI. Significance levels are as follows: * *p* ≤0.05, ** ≤0.01, *** ≤0.001, **** ≤0.0001, ***** ≤0.00001.

**Supplementary Tables**

**Table S1.** Clade-representative *Candida auris* strains used in this study: Representative strain for each major clade of *C. auris* used in the study, with notes on human disease source, country of discovery, and date of isolation. Mortality statistics are given in terms of absolute numbers of embryos over biological triplicate experiments. Additional significance testing is also noted for pairwise comparisons by Log-Rank test with Benjamini-Hochberg (BH) correction.

**Table S2:** Differentially expressed genes in *A. dispar* microinjection: All differentially expressed genes (DEGs) for *A. dispar* for (upper rows) sham injection *vs* no injection and (lower rows) *C. auris* injection *vs* sham injection. Log-fold change and -log_10_(false discovery rate) are calculated across all clades as a mean value for *C. auris* injection. Significant DEGs for each set of embryos injected with each clade are given (1 = significant, 0 = not significant), and the total number of clades where DEGs where significant is also given.

**Table S3.** Differentially expressed genes in *C. auris* infection: **(A)** Combined set of differentially expressed genes (DEGs) expressed by five clades during infection. Each set of DEGs expressed *in vivo vs in vitro* for five clades was combined, resulting in a set of genes that were specific to *C. auris* infection at these timepoints. **(B)** Combined set of DEGs between more virulent and less virulent clades: Each set of DEGs between clades I & IV (more virulent) and II & III (less virulent) was compared at 24 HPI and 48 HPI, resulting in a combined set expressed in infection. **(C)** Combined DEG Sets: between filamentous clade V and all other clades: Each set of DEGs between clade V and all each clades I-IV was compared at 24 HPI and 48 HPI, resulting in a combined set expressed in infection, excluding those significant during growth in YPD broth where clade V was not observed to filament. **(D)** Accessory genes not found in clade I reference genome: DEGs without orthologues in the clade I reference were identified and annotated. **(E)** Log-fold change and FDR across all pathogen gene expression. **(F)** Fragments per kilobase per million across all pathogen gene expression.

**Table S4.** Publicly available RNA-seq Data on 16^th^ April 2024: A systematic re-analysis of RNA expression across NCBI-available datasets for *C. auris* gene expression, categorised by research theme for analysis (**Figure S4****E-F**).
